# The Development of Active Binocular Vision under Normal and Alternate Rearing Conditions

**DOI:** 10.1101/2020.02.20.957449

**Authors:** Lukas Klimmasch, Johann Schneider, Alexander Lelais, Bertram E. Shi, Jochen Triesch

## Abstract

The development of binocular vision is an active learning process comprising the development of disparity tuned neurons in visual cortex and the establishment of precise vergence control of the eyes. We present a computational model for the learning and self-calibration of active binocular vision based on the Active Efficient Coding framework, an extension of classic efficient coding ideas to active perception. Under normal rearing conditions, the model develops disparity tuned neurons and precise vergence control, allowing it to correctly interpret random dot stereogramms. Under altered rearing conditions modeled after neurophysiological experiments, the model qualitatively reproduces key experimental findings on changes in binocularity and disparity tuning. Furthermore, the model makes testable predictions regarding how altered rearing conditions impede the learning of precise vergence control. Finally, the model predicts a surprising new effect that impaired vergence control affects the statistics of orientation tuning in visual cortical neurons.

## Introduction

Humans and other species learn to perceive the world largely autonomously. This is in sharp contrast to today’s machine learning approaches (***Kotsiantis et al., 2007***; ***Jordan and Mitchell, 2015***), which typically use millions of carefully labeled training images in order to learn to, say, recognize an object or perceive its three-dimensional structure. How can biological vision systems learn so much more autonomously? The development of binocular vision presents a paradigmatic case for studying this question. This development is an active process that includes the learning of appropriate sensory representations and the learning of precise motor behavior. Species with two forward facing eyes learn to register small differences between the images projected onto the left and right retinas. These differences are called binocular disparities and are detected by populations of neurons in visual cortex (***Kandel et al., 2000***; ***Blake and Wilson, 2011***) that have receptive subfields in both eyes. Frequently, they are modeled using separate Gabor-shaped filters for each eye, where the disparity is encoded by a shift in the centers of the filters, a difference between their phases, or by a combination of both (***Fleet et al., 1996***; ***Chen and Qian, 2004***). The responses of such disparity tuned neurons can be used to infer the three-dimensional structure of the world. At the same time, we also learn to align our eyes such that the optical axes of our two eyes converge on the same point of interest. These so-called vergence eye movements are also learned and fine-tuned during development (***Held et al., 1980***; ***Fox et al., 1980***; ***Stidwill and Fletcher, 2017***). Again, this learning does not require any obvious help from outside, but must rely on some form of self-calibration.

While it has long been argued that the development of disparity tuning and vergence eye movements are interdependent (***Hubel and Wiesel, 1965***), it has been only recently that computational models have tried to explain how the learning of disparity tuning and vergence eye movements are coupled and allow the visual system to self-calibrate (***Franz and Triesch, 2007***; ***Zhao et al., 2012***; ***Klimmasch et al., 2017***; ***Eckmann et al., 2019***). These models have been developed in the framework of Active Efficient Coding (AEC), which is an extension of Barlow’s classic efficient coding hypothesis to active perception (***Barlow, 1961***). In a nutshell, classic efficient coding argues that sensory systems should use representations that remove redundancies from sensory signals to encode them more efficiently. Therefore, sensory representations should be adapted to the statistics of sensory signals. Based on this idea, a wide range of data on tuning properties of sensory neurons in different modalities have been explained from a unified theoretical framework (***Dan et al., 1996***; ***Vinje and Gallant, 2000***; ***Simoncelli, 2003***; ***Smith and Lewicki, 2006***; ***Doi et al., 2012***). AEC goes beyond classic efficient coding by acknowledging that developing sensory systems shape the statistics of sensory signals through their own behavior. This gives them a second route for optimizing the encoding of sensory signals by adapting their behavior. In the case of binocular vision, for example, the control of vergence eye movements is shaping the statistics of binocular disparities. By simultaneously optimizing neural tuning properties and behavior, AEC models have provided the first comprehensive account of how humans and other binocular species may self-calibrate their binocular vision through the simultaneous learning of disparity tuning and vergence control.

A generic AEC model has two components. The first component is an efficient coding model that learns to encode sensory signals by adapting the tuning properties of a population of simulated sensory neurons (***Olshausen et al., 1996***; ***Olshausen and Field, 1997***). In the case of binocular vision, this is a population of visual cortical neurons receiving input from the two eyes that learns to encode the visual signals via an efficient code. The second component is a reinforcement learning (RL) model that learns to control the behavior. In the case of binocular vision, this component will learn to control eye vergence. For this, it receives as input the population activity of the visual neurons and learns to map it onto vergence commands. This learning is guided by an internally generated reward signal, which reinforces movements that lead to a more efficient encoding of the current visual scene. For example, when the eyes are aligned on the same point, the left and right images become largely redundant. The efficient coding model can exploit this redundant structure in both eyes, by developing neurons tuned to small or zero disparities. Conversely, such binocular neurons tuned to small disparities will represent any remaining misalignments of the eyes, providing informative input for vergence control. In this way, learning of vergence control supports the development of neurons tuned to small disparities and this developing population of neurons in turn facilitates the learning of fine vergence control (***Zhao et al., 2012***).

Importantly, however, this normal development of binocular vision is impaired in a range of alternate rearing conditions. In fact, already since the days of Hubel and Wiesel, alternate rearing conditions have been used to improve our understanding of visual cortex plasticity and function. Manipulating the input to the visual system during development and observing how the system reacts to such manipulations has shaped our understanding of visual development until today. For example, artificially inducing a strabismus leads to drastic changes in the tuning properties of neurons in visual cortex (***Hubel and Wiesel, 1965***). A comprehensive theoretical account of the development of binocular vision must therefore also be able to explain the experimentally observed differences in alternate rearing conditions. Therefore, we here test if a recently proposed AEC model of the development of binocular vision can reproduce and explain the large range of experimental findings from different alternate rearing conditions. Indeed, we show that the model qualitatively captures findings on how different alternate rearing conditions alter the statistics of disparity tuning and binocularity. Furthermore, the model makes specific novel and testable predictions about differences in vergence behavior under the different rearing conditions. Surprisingly, it also predicts systematic differences in the statistics of orientation tuning of visual cortical neurons depending on the fidelity of vergence eye movements. Overall, our results support AEC as a parsimonious account of the development of binocular vision, highlighting the active nature of this development.

## Results

### A model for the development of active binocular vision

The model comprises a virtual agent situated in a simulated environment. The agent looks at a textured plane, on which images from the man-made section of the McGill Database (***Olmos and Kingdom, 2004a***) are rendered. The plane is positioned in front the agent at variable distances (Fig. 1A). An image is rendered for the left eye and a second image is rendered for the right eye. Binocular patches are extracted from these images and encoded by a sparse coding algorithm. The activation levels of the learned binocular basis functions (BFs) can be thought of as firing rates of binocular simple cells in primary visual cortex. The basis functions themselves roughly describe their receptive fields and are optimized through learning (***Olshausen and Field, 1997***). These activations are then squared and pooled across the image to obtain a more position-invariant representations mimicking the behavior of complex cells. From this state representation a reinforcement learner generates vergence commands that symmetrically rotate the eyeballs inwards our outwards. This results in two new images being rendered and a new simulation iteration starts. The complete process is depicted in Fig. 1B (see Methods for details).

**Figure 1.**
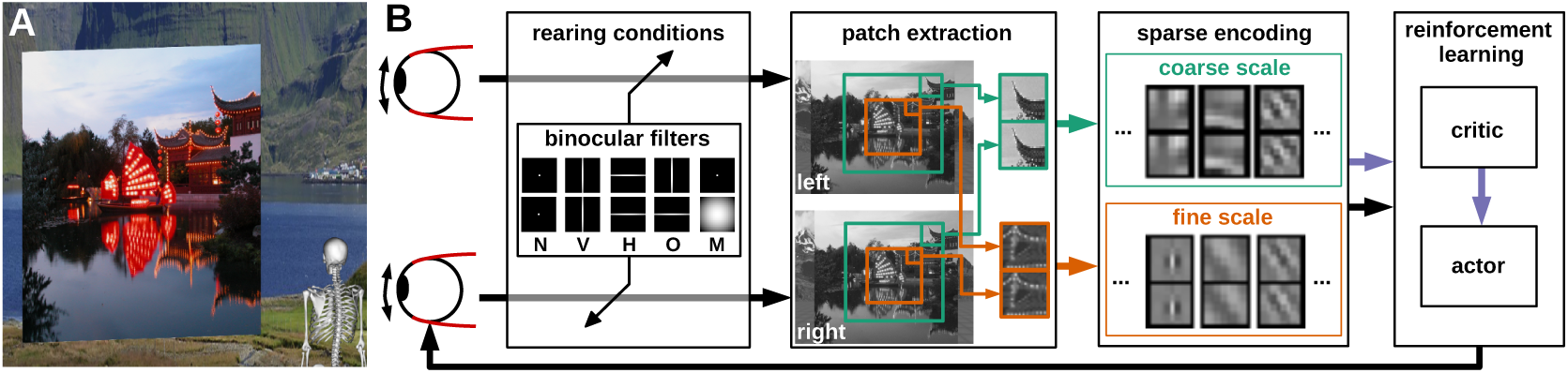
Model overview. **A** The agent looking at the object plane in the simulation environment. **B** Processing steps of the active efficient coding model. One image is generated per eye. We convolve them with different kernels, such as those in the inbox, to simulate alternate rearing conditions (N: normal, V: vertical, H: horizontal, O: orthogonal, M: monocular). Binocular patches are extracted in a coarse and a fine scale (green and orange boxes) with different resolutions. These patches are encoded by activations of basis functions via sparse coding and combined with the muscle activations to generate a state vector. While this vector is fed into the reinforcement learning architecture, the sparse coding step also generates a reconstruction error that indicates the efficiency of encoding. We use this signal as reward (purple arrow) to train the critic, which in turn evaluates states to teach the actor. Finally, the actor generates changes in muscle activations, which result in rotations of the eyeballs and a new iteration of the perception-action cycle.

In the human retina, the receptive field (RF) size of ganglion cells increases towards the periphery (***Curcio et al., 1990***). We incorporate this idea by extracting patches from an input image at two different spacial scales: A high-resolution fine scale is extracted from the central part and a low-resolution coarse scale is extracted from a larger area (orange and turquoise boxes in Fig. 1 and 2). Covering a visual angle of 8.3° in total, the fine scale corresponds to the central/para-central region (including the fovea) and the coarse scale to the near-peripheral region with a diameter of 26.6°. On the one hand, this two-scale architecture is more biologically plausible than using just a single scale, on the other hand it also increases the resulting verging performance (***Lonini et al., 2013***). One input patch (or subfield) in the coarse scale can detect a disparity of up to 8.8° while one patch in the fine scale covers 1.6°. The coarse scale can therefore be used to detect big disparities, while the fine scale represents small disparities.

**Figure 2.**
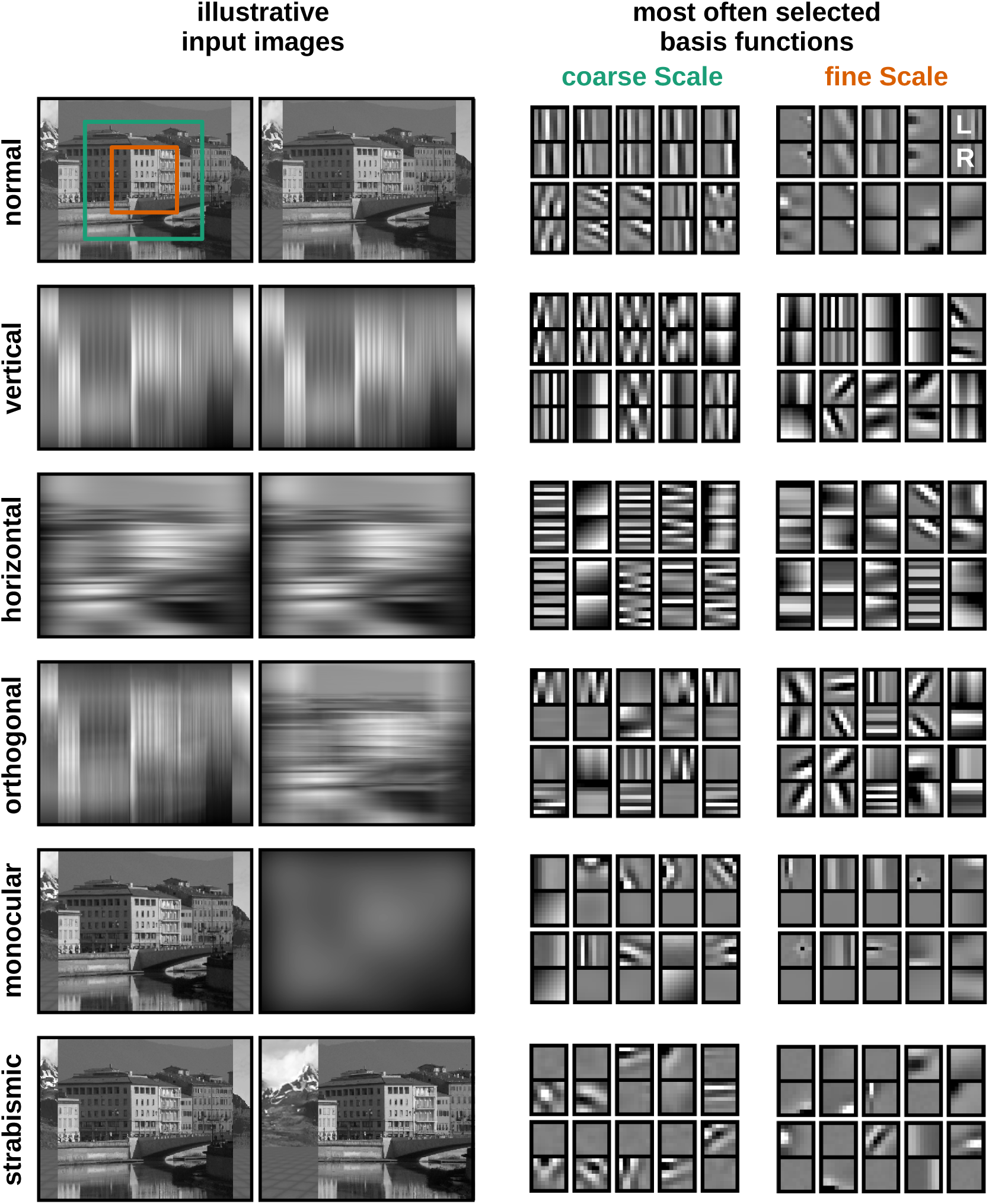
Input scenarios and learned receptive fields. **Left:** Illustration of the input under different rearing conditions. Except for the normal scenario, the images are convolved with different Gaussian filters to blur out certain orientations or simulate monocular deprivation. To simulate strabism the right eye is rotated inward by 10°, so that binocular neurons receive non-corresponding inputs to their left and right eye receptive fields. **Right:** Representative examples of binocular basis functions (BFs) for the fine and coarse scale learned under the different rearing conditions after 0.5 million iterations. For each BF the left eye and right eye patch are aligned vertically. In each case, the 10 BFs selected most frequently by the sparse coding algorithm are shown.

We simulate altered rearing conditions by convolving the input images for the two eyes with two-dimensional Gaussian kernels to blur certain oriented edges, or to simulate monocular deprivation. To mimic strabismus, the right eyeball is rotated inwards while the left eye remains unchanged to enforce non-overlapping input to corresponding positions of the left and right retina (see Methods for details).

The adaptation of the neural representation and the learning of appropriate motor commands occur simultaneously: While the sparse coder updates the BFs to minimize the reconstruction error, the RL agent generates vergence eye movements to minimize the reconstruction error of the sparse coder. Since the sparse coder has a fixed capacity, minimizing its reconstruction error is equivalent to maximizing its coding efficiency. Thus, both the sparse coder and the reinforcement learner aim to maximize the overall coding efficiency of the model.

### Normal rearing conditions lead to the autonomous learning of accurate vergence control for natural input and random dot stereograms

Under normal rearing conditions the joint learning of the neural representation and motor behavior results in an agent that accurately verges the eyes on the plane in front of it (***Klimmasch et al., 2017***). This behavior emerges in an autonomous fashion, since both the sparse coder and the RL agent only strive to improve the neural encoding by reducing the reconstruction error. We demonstrate this behavior in Video 1 (videos/vergence_movements.mp4) and will analyse it in greater detail in the following sections.

A critical test of any model of the development of stereoscopic vision is whether it can handle *random-dot stereograms* (RDSs), which represent the most challenging stimuli for stereopsis. Since their introduction by ***Julesz (1971***) RDSs have been used extensively to investigate the human ability for stereoscopic vision. Nowadays they are used in opthalmological examinations to asses stereo acuity as well as to detect malfunctions in the visual system, such as strabismus or amblyopia (***Walraven, 1975***; ***Okuda et al., 1977***; ***Ruttum, 1988***). In these experiments participants view a grid of random dots through a stereoscope or another form of dichoptic presentation. Typically, the central part is shifted in one of the two images which results in the perception of stereoscopic depth in healthy subjects. The advantage of this form of examination is that there are no monocular depth cues (such as occlusion, relative size, or perspective). The impression of depths arises solely because of the brain’s ability to integrate information coming from the two eyes.

To show that our model is able to perceive depth in RDS, we generate various RDS and render the shifted images for the left and right eye separately. We expose the model that was trained on natural input stimuli to a range of RDS with different spatial frequencies, window sizes, disparities, and object distances. A video of the performance can be found in Video 2 (videos/performance_on_RDS.mp4). The model is clearly able to detect the differences in the images and align the eyes on the virtual plane that will appear in front or behind the actual object plane in the RDS. Averaged over all trials, the model achieves a vergence error of 0.2°. This is comparable with our results on natural images (see Fig. 6) and indicates that the model generalizes well to artificial images it has never seen before.

### Altered rearing conditions cause changes in neural representation matching experimental findings

A second critical test of any model of the development of binocular vision is whether it can account for the effects of alternate rearing conditions observed in biological experiments. We simulate such alternate rearing conditions by filtering the input images for the left and right eyes with Gaussian filters. Figure 2 shows illustrative examples of the filtered images that were used to train our model and the respective learned BFs.

When the model is trained with unaltered natural visual input, the resulting RFs resemble Gabor wavelets (***Daugman, 1985***), as shown in the first row in Fig. 2. They appear similar in the coarse and the fine scale, but tend to be more localized in the latter. The changes that are applied to the input images in the alternate rearing conditions are reflected in the RFs that are learned: Among the 10 most often selected BFs there are no vertically (horizontally) oriented RFs, when the model is trained on images that are deprived of vertical (horizontal) edges. Orthogonal RFs emerge as a result of training on orthogonal input. When one eye is deprived of input, the RFs will become *monocular* and encode information coming from the “healthy” eye only. Strabismic rearing results in the development of monocular RFs without a preference for one or the other eye (***Hunt et al., 2013***).

The full set of all BFs (coarse and fine scale) for all the rearing conditions can be found in supplemental Fig. 1. The following sections will analyse them in a more quantitative fashion.

### Neurons’ orientation tuning reflects input statistics

To analyze the statistics of the developing RFs in greater detail, we fit oriented two-dimensional Gabor wavelets to each BF. For this part of the analysis the left and right parts of the binocular BFs are studied separately, so we look at the *monocular* BF fits only. We combine the results from coarse and fine scale, since a two-sample Kolmogorov-Smirnov test (***Young, 1977***) did not reveal a statistically significant difference between the distributions. Only those BFs which met a criterion for a sufficiently good fit (98% of all bases) are considered for further analysis (see Methods).

Figure 3 shows how the altered input changes the distribution of preferred orientations of the BFs. In the *normal* case we can observe a clear over-representation of vertically (0°) and horizontally (90°) tuned BFs. This is known as the *oblique effect* and has been frequently observed in animals (***Appelle, 1972***; ***Li et al., 2003***) and humans (***Furmanski and Engel, 2000***). It has been argued that it stems from the over-representation of vertical and horizontal edges in natural images (***Coppola et al., 1998***). Additionally, we cannot exclude the possibility that it is related to the rectangular pixel grid representing the input to our model.

**Figure 3.**
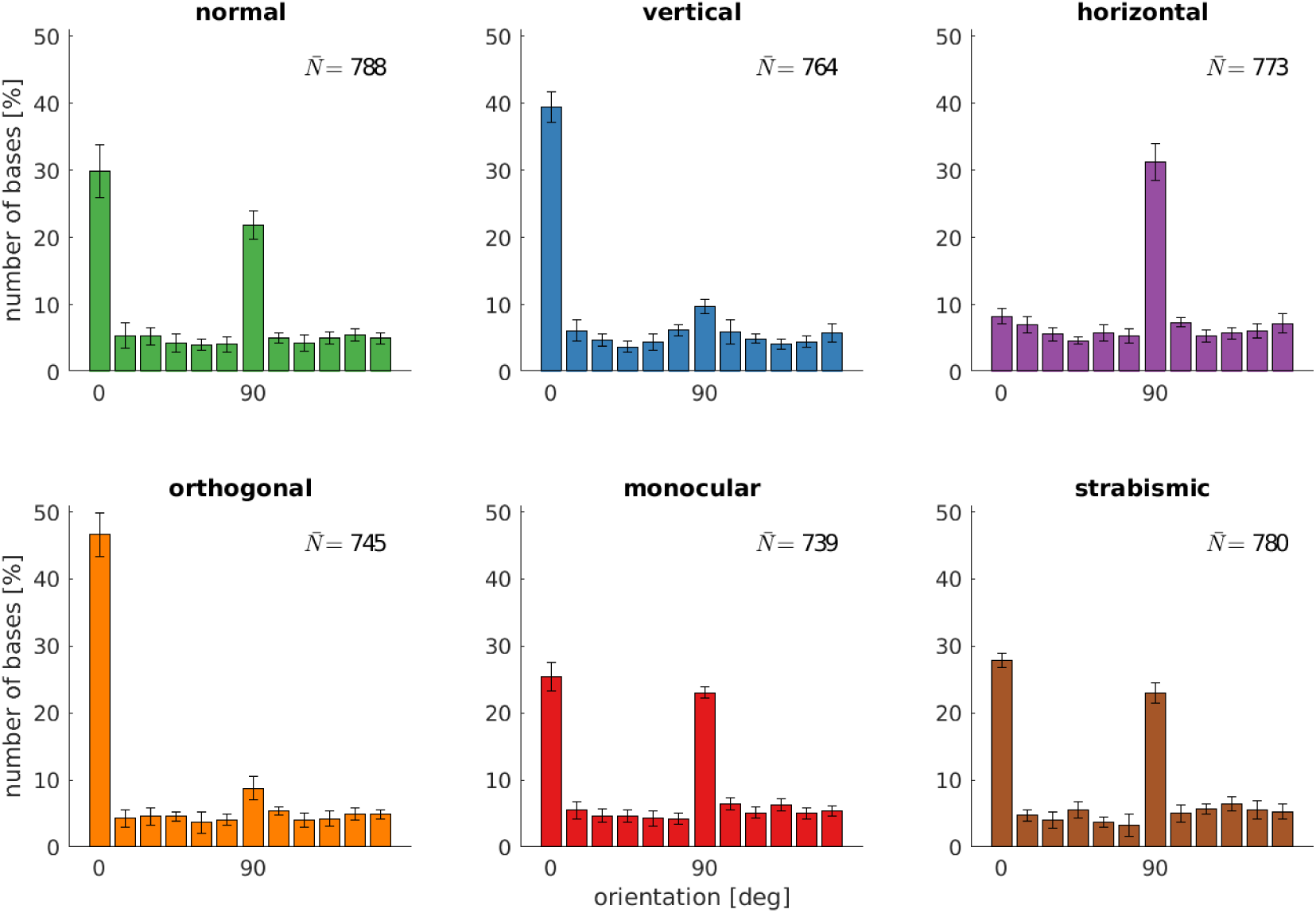
Orientation distributions for different rearing conditions. Displayed are the orientations of the Gabor wavelets that were fitted to the learned BFs of the left eye. Shown are the best fits from coarse and fine scale combined (800 in total). The error bars indicate the standard deviation over 5 different simulations. 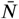 describes the average number of BFs that passed the selection criterion for their fits (see Methods).

While the distribution of orientations does not change much in the *monocular* and *strabismic* rearing case, we observe a marked difference to the normal case when certain orientations are attenuated in the input. The models trained on *vertical* input are missing the peak at horizontal orientations and vice versa for the *horizontal* case. Additionally, we find an increased number of neurons tuned to the dominant orientation in the input. These observations are consistent with animal studies (***Stryker et al., 1978***; ***Tanaka et al., 2006***)

The separate analysis of the RFs in the left and right eye for the models that were trained on *orthogonal* input reveals the adaptation of each eye to its input statistics. Furthermore, we find that orthogonal RFs developed (also see fourth row in Fig. 2) that have been observed in an orthogonal rearing study in cats (***Leventhal and Hirsch, 1975***).

### The development of binocular receptive fields requires congruent input to the two eyes

Another interesting feature of the neural representation that has been studied extensively in the context of alternate rearing is the *binocularity*. The binocularity index (BI) is used to assess how *responsive* a neuron is to the inputs from the two eyes. A *binocular* neuron requires input from both eyes to respond maximally, while a *monocular* neuron is mostly driven by just one eye.

To determine the binocularity indices for the neurons in our model we use the original method from ***Hubel and Wiesel (1962***). They determined a stimulus that maximizes the monocular response, and applying this stimulus separately in left or right eye to get the (monocular) neural responses *L* and *R* (see Methods for details). For each neuron the binocularity index is then calculated as 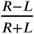. Like Hubel and Wiesel we sort the binocularity indices into seven bins. The values range from -1 (monocular left) over 0 (binocular) to +1 (monocular right).

Figure 4 depicts the binocularity distributions for the coarse and the fine scale for all rearing conditions. The models that were trained on input that is coherent between the left and right eye (top row) exhibit the majority of neurons falling in the bin with binocularity index 0. Neurons in this category receive about the same drive from the left and the right eye. In the *normal* case more neurons fall into that bin than in the *vertical* and *horizontal* case. This is due to the ability of the model to perform precise vergence control: Since left and right image are almost identical most of the time, the great majority of basis functions will develop to encode the exact same input from both eyes. This, in turn, will result in the cells being completely binocular with a binocularity index of 0. In the vertical and horizontal case, we observe a reduction in the number of cells that have a binocularity index of 0. We attribute this to the limited vergence performance in these cases, that we will analyse in the next sections.

**Figure 4.**
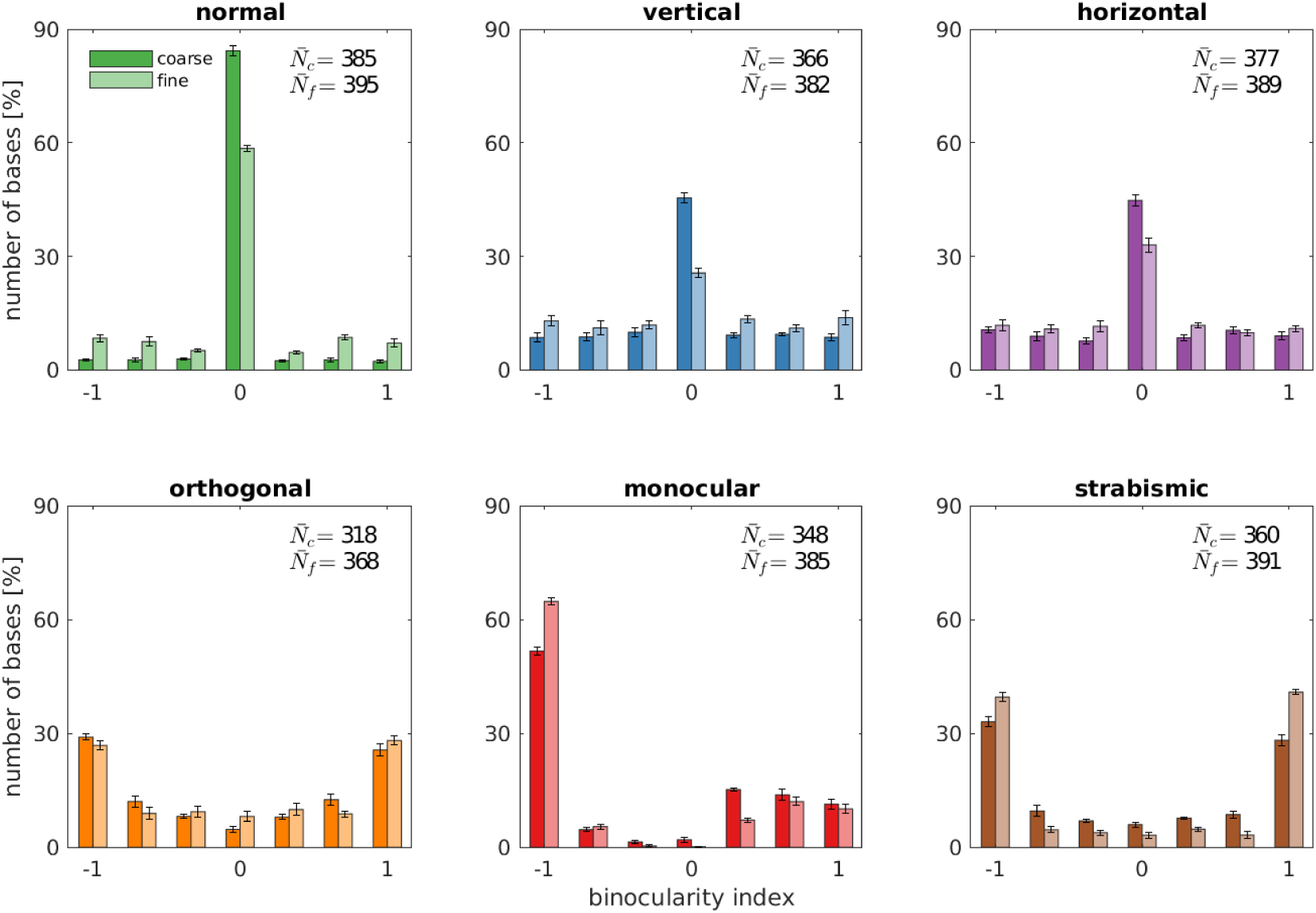
Binocularity distributions for different rearing conditions. The binocularity index is calculated by comparing the neuron’s responses to monocular stimuli. The values range from -1 (monocular left) over 0 (binocular) to 1 (monocular right). Results for coarse and fine scale are presented next to each other. Error bars indicate the standard deviation over 5 different simulations. 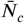 and 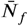 are the average number of basis functions (out of a total of 400) that pass the selection criteria for their fits (see Methods).

If, on the other hand, the input differs qualitatively for the two eyes (Fig. 4, bottom row) the receptive fields will also differ in their monocular sub-parts. This can also be observed in Fig. 2 for the orthogonal, monocular and strabismic case. Looking at the binocularity index, we find that most of the cells become monocular, with a symmetric distribution for orthogonal and strabismic rearing. Monocular deprivation of the right eye leads to a distribution of binocularity indices that is mostly monocular for the left eye.

We also find differences between coarse and fine scale, with slightly fewer binocular and slightly more monocular cells in the latter one. This indicates that left and right part of the BFs in the fine scale tend to be marginally more different than in the coarse scale. Patches that serve as input to this scale are not down-sampled and have a high resolution. Small differences in the input patches will therefore not be blurred out and lead to small differences in the learned BFs since the sparse coder strives for reconstructing the input as accurately as possible.

Looking into the biological data, we find the pronounced peak at binocular neurons in the normal case (***Wiesel and Hubel (1963***), Fig. 1, and ***Hubel and Wiesel (1965***), Fig. 5). When trained on inputs deprived of certain orientations (***Stryker et al. (1978***), Fig. 6B), the neurons become relatively more monocular, but most of the neurons remain binocular. This is in good agreement with our model.

**Figure 5.**
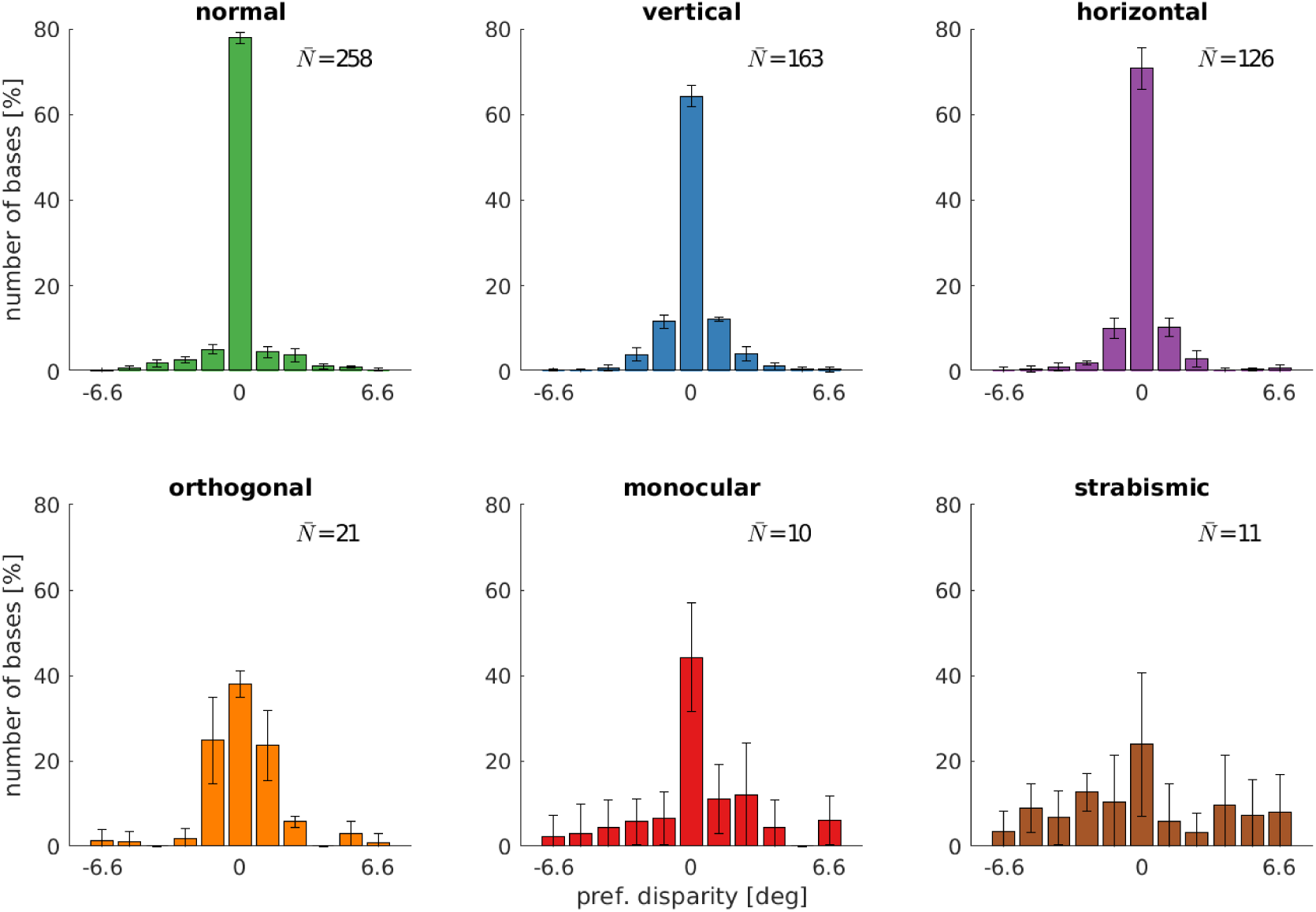
Disparity distributions for different rearing conditions. The neuron’s preferred disparities are extracted from the binocular Gabor fits. All neurons with a disparity bigger than the maximally detectable one are removed from the analysis. Presented are the averaged data of the coarse scale from 5 random seeds. 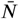 describes the average number of neurons that met the selection criteria (see Methods).

**Figure 6.**
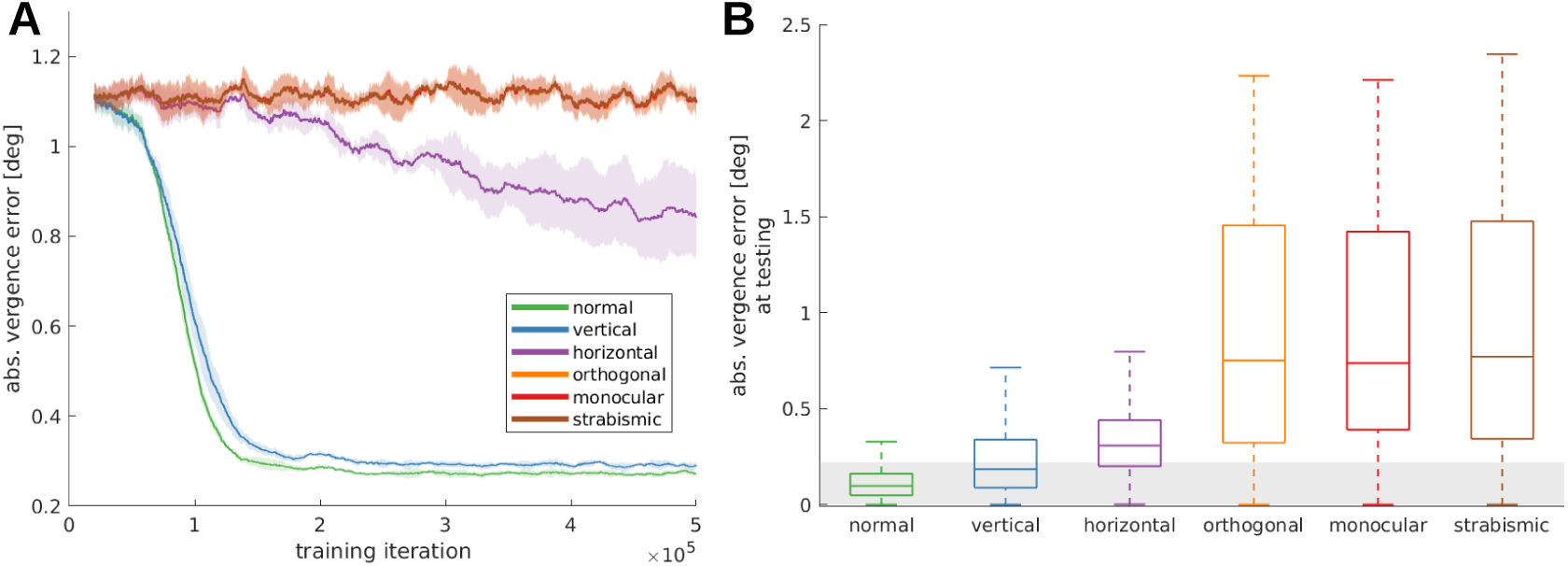
Vergence performance of models raised under different rearing conditions. **A** Moving average of the vergence error over the duration of the training period. During the training, a textured object plane is positioned in front of the agent at varying distances. The vergence error is defined as the difference between the angle that is desired to fixate exactly on the object plane and the actual angle between the eyes. The shadows indicate the standard deviation over 5 different random seeds. **B** Vergence errors after 20 perception-action-cycles on unknown input stimuli starting from various initial errors. This testing was done without the visual aberrations encountered during training. Displayed are conventional box plots without outliers. The gray bar indicates a vergence error of 0.2° which presents the resolution boundary of our system.

***Stryker et al. (1978***) reared kittens on orthogonal input and report an increase in monocular neurons (Fig. 6A) when compared to the normal rearing data from Hubel and Wiesel. In comparison to the rearing on stripes, there are fewer binocular cells. The loss of binocular neurons that we see in our data is also reported in ***Hirsch and Spinelli (1970***), who reared kittens on orthogonal stripes. Monocular rearing and the analysis of binocularity was performed in ***Wiesel and Hubel (1963***). In Fig. 3 and 5 we see the development of completely monocular cells after visual deprivation of the other eye. The strabismic case was studied a few years later in ***Hubel and Wiesel (1965***) (Fig. 5A) and revealed a division of the neural population in monocular neurons for either left or right eye, in agreement with our model.

### Alternate rearing conditions reduce the number of disparity tuned cells

A central aspect of the development of binocular vision is the emergence of neurons which are tuned to binocular disparities. We therefore investigate how alternate rearing affects the number of neurons with disparity tuning in the model and the distribution of their preferred disparities. We estimate disparity tuning by considering phase shifts between left and right RFs in the following way: We fit binocular Gabor wavelets to the BFs, where all parameters, except for the phase shift, are enforced to be identical for the left and right monocular BF. The disparity for one neuron can then be calculated as described in Analysis of receptive fields. The distribution of disparity tuning of the coarse scale neurons is shown in Fig. 5 for the different rearing conditions. Results for the fine scale are comparable and presented in supplemental Fig. 2. First, there is a noticeable difference in the number of cells that are disparity tuned between the different rearing conditions: In the normal case we find the highest number of disparity tuned cells, rearing in a striped environment reduces the number, and uncorrelated input results in the smallest number of disparity tuned cells. In every case, the distribution of preferred disparities is peaked at zero. The height of this peak is reduced for rearing conditions with in-congruent input to the two eyes.

Comparing the normal with the vertical and horizontal case, there is an increase in the number of cells that are tuned to non-zero disparities. This indicates that under these alternate rearing condition, the agents are exposed to non-zero disparities more often. This is in good agreement with the results from the next section (also see Fig. 6), where we will see that those models perform less accurate vergence movements compared to the normal case.

In the strabismic case, a neuron’s receptive fields in left and right eye are driven by un-corresponding input. This results in very few disparity tuned cells that exhibit a much broader distribution of preferred disparities.

To investigate the effect of a less severe strabism we conduct an additional experiment similar to ***Shlaer (1971***) (see Fig. 2). Here, we fix the strabismic angle to 3°, which results in a corresponding image in the two eyes because one input patch in the coarse scale covers an angle of 6.4°. *Supplemental Fig. 3* shows that this leads to an increased amount of disparity tuned cells and a shift of their preferred disparity to 3°. Exactly as in ***Shlaer (1971***), the constant exposure to a certain disparity leads to a preference to that disparity for the majority of cells.

### Model predicts how alternate rearing conditions affect vergence learning

While the effect of alternate rearing conditions on receptive fields of visual cortical neurons is well studied, there has been little research on the effect of alternate rearing conditions on vergence behavior. To quantify vergence behavior in the model, we define the absolute vergence error. It measures by how much the vergence angle between the eyes deviates from the ideal position, which would make the two eyes fixate the same center of the object. This measurement is taken at the end of a fixation (corresponding to the last of 10 time steps), to give the model sufficient time to fixate the object.

Figure 6A shows the evolution of the absolute vergence error over training time for the different rearing conditions. The models with *normal* or *vertical* rearing learn to verge the eyes on the same point on the object, resulting in the reduction of the vergence error to small values of around 0.3 degrees. The model that learns on images without vertical edges (*horizontal* case) does manage to verge the eyes slightly, but does not reach the accuracy of the former models. The *orthogonally, monocularly* and *strabismically* reared models do not improve much in comparison to random behavior in the beginning of training. The main difference to the models that were able to learn vergence is that under these conditions the left and right eye are provided with in-congruent input. The orthogonal model receives two monocular images that retain different orientations. The right monocular image of the monocularly deprived model contains little information at all, and the two eyes are physically prevented from looking at the same object in the strabismic case. In these cases, very few neurons with disparity tuning emerge (compare previous section) that could drive accurate vergence eye movements.

### Behavior after normal visual input is reinstated

Alterations of the visual input during the critical period of visual development lead to lasting visual deficits. To simulate the effect of a transient alteration of visual input during the critical period, we first train the model under alternate rearing conditions as described above and then reinstate normal visual input. For this, we freeze all weights after the training phase and test all models on the same, un-altered input images. By doing so, we simulate a situation where the visual aberrations present during development (such as astigmatism or a cataract) are corrected *after* the critical period.

The object plane is put to a distance ∈ [0.5, 1, …, 6] m, the initial vergence error is chosen randomly from −2 to 2°, and 40 stimuli that were not seen during training are applied on the object plane. From these initial conditions we run the simulation for 20 iterations and record the vergence error at the end of each fixation. The results of this testing procedure are displayed in Fig. 6B. Here, the gray shaded area indicates a vergence error of 1 pixel. We observe that the *normally* trained model exhibits the best performance and actually achieves sub-pixel accuracy in the great majority of trials. Interestingly, the performance declines for the *vertical* model. One could expect the model that was trained solely on vertical edges to be better at aligning those edges. We attribute this to mis-alignments (or false matches) between the two images that happen more frequently, when the world is made up only of vertical edges. Additionally, the neural representation that was learned during the exposure to vertical edges only might not be utilized as efficiently as before, now that all orientations are present in the input.

Even though the performance of the model trained on only *horizontal* orientations is quite poor during training, after applying the correction it clearly displays a verging behavior. In comparison to the *orthogonal, monocular* and *strabismic* models, it reduces the vergence error, though being less accurate than the other two cases.

The main difference between the conditions under which vergence could or could not be learned is the correspondence between the input images. When the inputs to the two eyes are in-congruent — as in the orthogonal, monocular and strabismic cases — we could not observe any improvement in the vergence error. Matching input, on the other hand, always led to the learning of vergence behavior. This becomes apparent especially after testing the learned models on un-altered inputs.

Since this is the first study to investigate the quality of learned vergence movements after exposure to alternate rearing conditions (to the best of our knowledge), the differences in performance are a genuine prediction of our model.

To explain this phenomenon we consider the reward function, which shapes the model’s behavior during training. Figure 7 shows the averaged reconstruction error over three different object distances and ten stimuli for the different rearing conditions. We defined the reward as the negative reconstruction error of the sparse coders. In the normal case, we clearly see an optimum of the reconstruction error at zero disparity. This also holds for the vertical and horizontal condition, whereas those are at least one magnitude smaller. We argue that the differences in the rewards lead to the differences in vergence performance, since all models that could not verge display a reward function that is rather flat for different disparity values. The models with a negative peak at zero disparity, on the other hand, all learn to verge and the difference in the magnitude of the reward seems to be reflected in the vergence performance after training.

**Figure 7.**
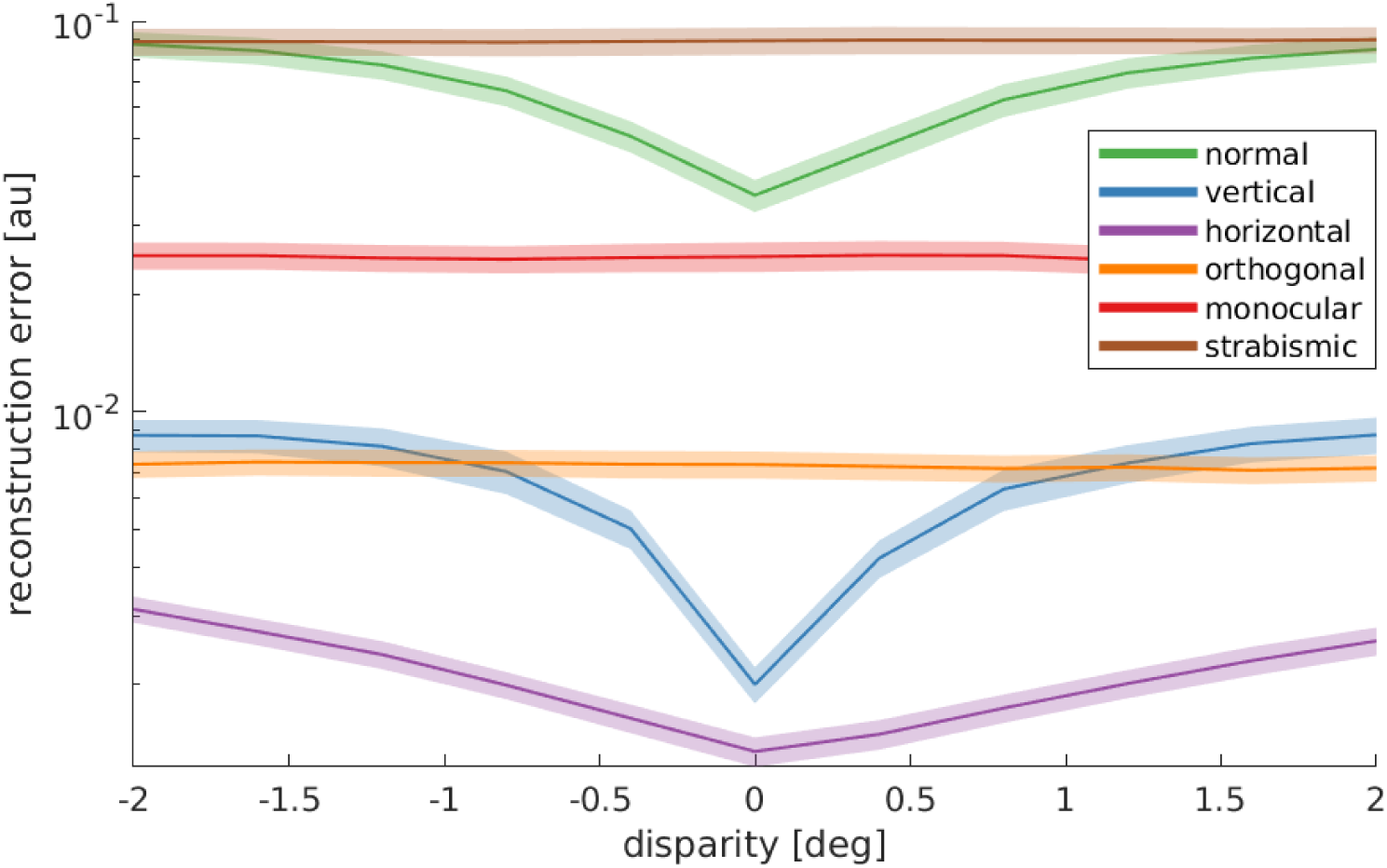
Reward functions for the different rearing conditions. The reward function is what drives the reinforcement learner to move the eyes in a useful fashion. For all different conditions, we plot the rewards that the models will receive at different disparities. Notice the log-scale on the y-axis. The data are averaged over 10 stimuli that were not encountered during training, three different object distances (0.5, 3, and 6m), and 5 random seeds for every condition. The shaded area represents the standard error. Only those models that receive corresponding input in left and right eye display a reconstruction error that is minimal at zero disparity. These are the only models that learn to verge the eyes.

### Model predicts how vergence movements influence the statistics of orientation preference

Our model also allows us to investigate, for the first time, how the quality of the vergence control influences the neural representation. As a baseline, we consider the orientation tuning of a reference model which was trained on normal images and learned an appropriate vergence policy. We compare this model to a version that was trained on the same input images, but could not verge the eyes. Specifically, the sparse coder was trained normally, but the RL part was removed. This model saw different disparities during training by looking at objects at different depths, but was not able to change this distribution of disparities to facilitate the encoding. We refer to this model as the “random disparity” model. In another version of the model, we artificially always set the vergence angle to correctly fixate the objects. In this way, this model was never exposed to disparities (except for very small ones in the periphery that arise because of slightly different perspectives in the left and right eye). We refer to this version as the “zero disparity” model.

Figure 8A shows the fraction of neurons that are tuned to vertical orientations (0± 15°) for these three models. When the influence of the RL agent is removed, we observe a significant decrease in the number of vertically tuned neurons. This change must be caused by the different distributions of disparities that the models experience due to their different motor behavior. In the model that was trained without disparities, we find the least amount of neurons tuned to vertical edges.

**Figure 8.**
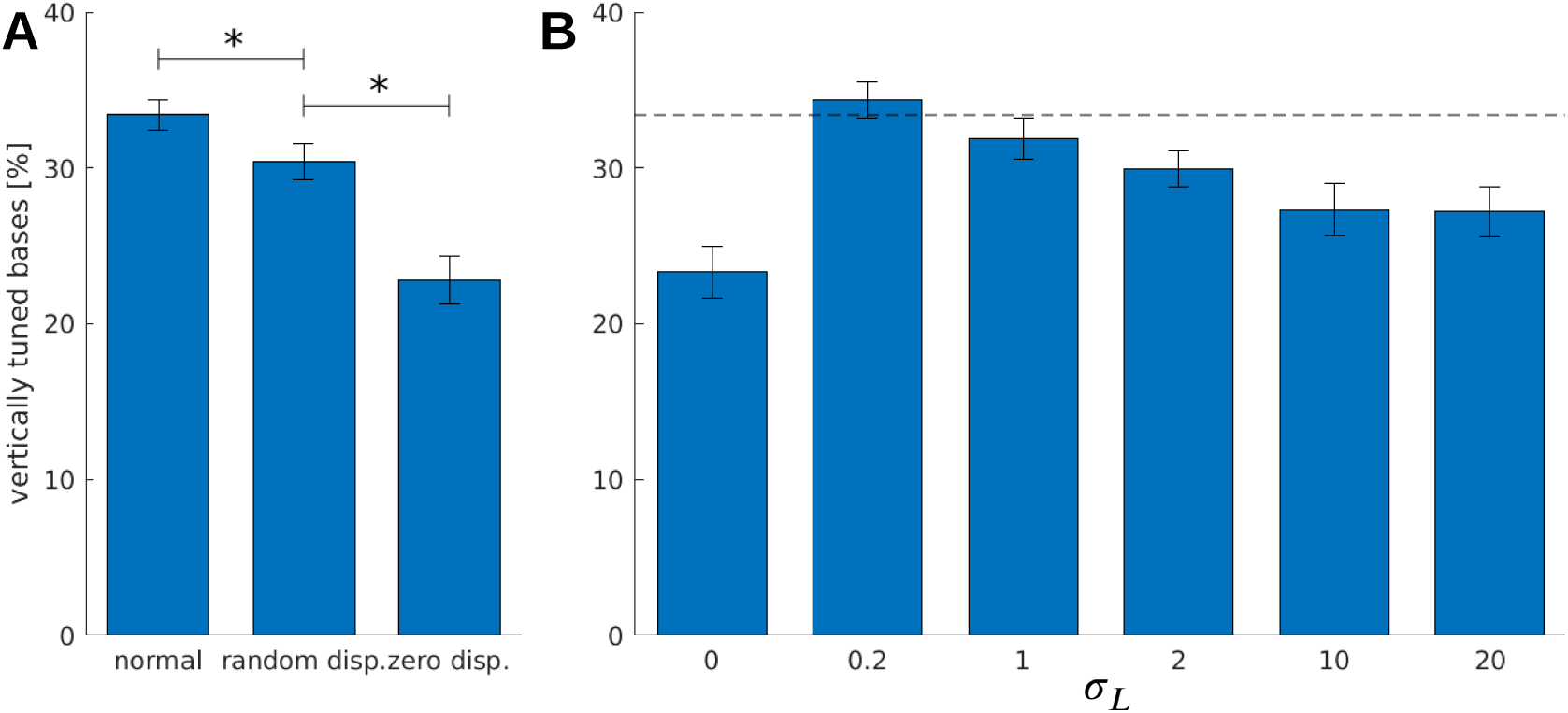
The effect on the learning of vergence and disparitites on the number of neurons tuned to vertical edges. **A** Here we compare the relative amount of cells that are tuned to vertical orientations for three different types of models: The first is the normal model, where the sparse coder and the RL agent are trained. In the second case, only the sparse coder is trained while the RL agent is removed. During the training procedure, this model is exposed to random disparities in the input images. In the last case, only the sparse coder is trained, but additionally, we artificially set the eyes to always verge perfectly on the objects in front of it. Like that, this model does not learn vergence movements and is not exposed to disparities as well. Asteriks indicate a statistically significant difference between the samples as revealed by the students t-test (p-values are 0.007 and 0.001). **B** These models were trained with a Laplacian distribution of different disparities. Depicted are the relative amount of BFs tuned to vertical orientations in dependence of *σ*_*L*_, the standard deviation of the Laplacian. *σ*_*L*_ =0 corresponds to 0 disparity all the time, while *σ*_*L*_ = 20 is an almost uniform disparity distribution. Error bars indicate the standard deviation over 5 different seeds. The black dotted line indicates the amount of vertically tuned neurons in the *normal* model.

To study the role of the distribution of experienced disparities more systematically, we train the sparse coder on different truncated Laplacian distributions of disparities. The distributions are heavy-tailed and centered around zero. The spread in this distribution is determined by the standard deviation *σ*_*L*_· *σ*_*L*_ =0 means zero disparity all the time (corresponding to the zero disparity case), while the distribution becomes almost uniform for big values of *σ*_*L*_· Figure 8B shows how the number of vertically tuned neurons changes in response to different values of *σ*_*L*_· We find the smallest number of vertically tuned cells when the disparity is zero throughout the whole training. For very large *σ*_*L*_ there are more vertical cells, but not as many as for smaller values which are different from zero. In fact for *σ*_*L*_ = 0.2, which corresponds to a standard deviation of one pixel in the input image, the number of vertically tuned neurons is maximized.

An intuitive explanation for this over-representation of cells tuned to vertical orientations is given in Fig. 9. Here, we depict a part of an input image at three different disparities. While the horizontal edge can be encoded by the same BF for all disparity values, the vertical edge demands three different basis functions to represent the input pattern faithfully. A system that experiences these disparities in its inputs, needs to devote neural resources to represent them all. If the distribution of disparities becomes too wide, however, individual neurons will receive close to independent input from both eyes and disparities that lie in the range that can be represented by a single basis function will be rare.

**Figure 9.**
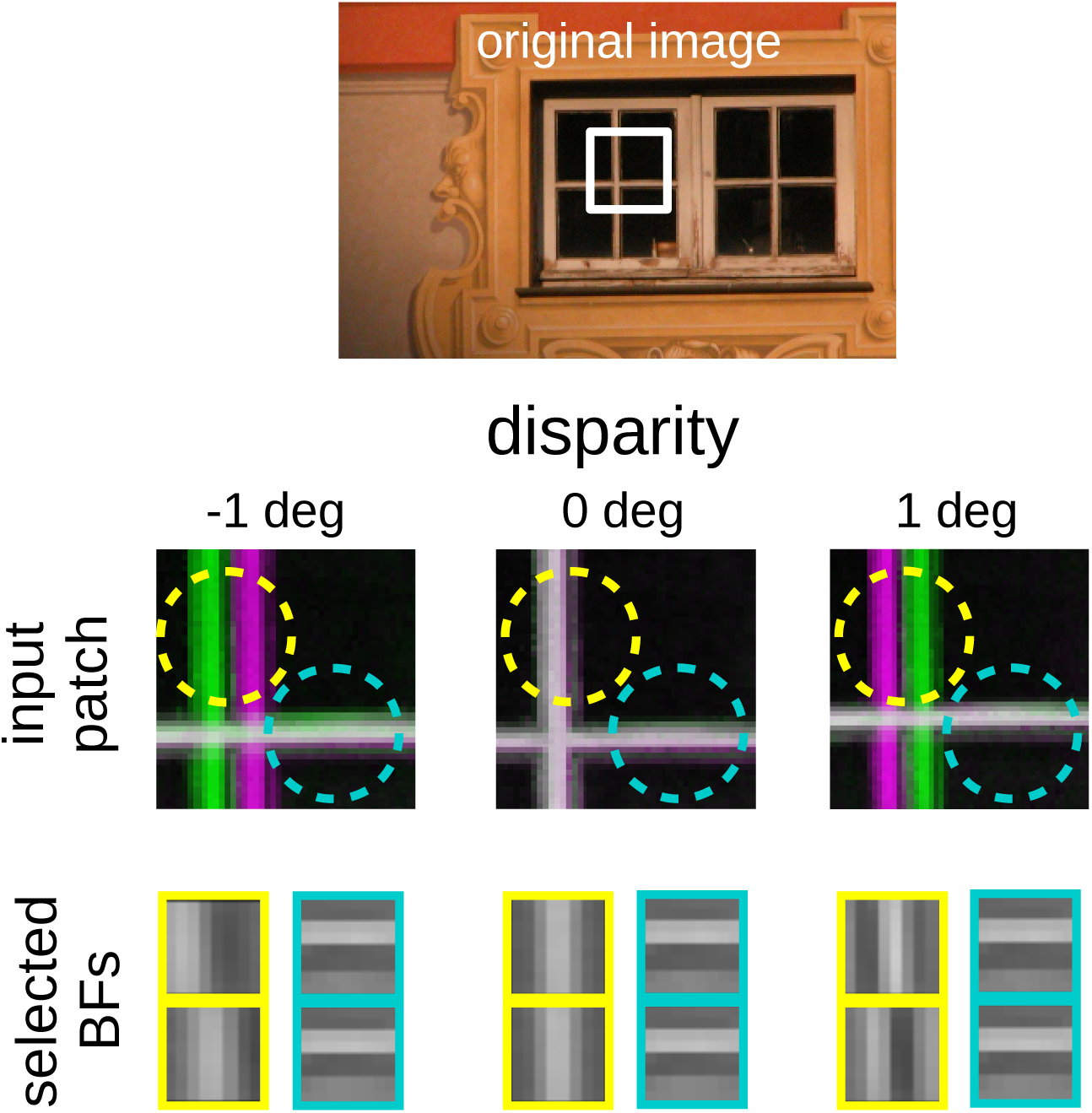
Intuition for the over-representation of vertical edges when disparities have to be encoded. We show the location of two RFs (yellow and cyan circles) on a patch in the visual field and present them with three different disparities. The inputs are depicted as anaglyphs, compositions of two images where the left image goes into the green channel and the right into a magenta channel. When the two images are corresponding, the anaglyph will appear in black and white, while un-corresponding parts will appear in green and magenta. For each disparity and RF we show the BF that is selected by the sparse coder to encode the input. While the BF that encodes the input in the cyan RF is the same for all disparities, the input inside the yellow RF can best be reconstructed by BFs that are tuned to that exact disparity.

## Discussion

A major goal of Computational Neuroscience is the development of models that explain how the tuning properties of neurons develop and how they contribute to the behavior of the organism. Over the last decades, the dominant theoretical framework for understanding the development of tuning properties of sensory neurons has been the *efficient coding hypothesis*. It states that sensory tuning properties adapt to the statistics of the sensory signals. In this framework, the behavior of the organism has been largely neglected, however. Specifically, there has been hardly any work on how developing neural tuning properties shape behavior, how the developing behavior affects the statistics of sensory signals, and how these changing statistics feed back on neural tuning properties. We argue that understanding the development of sensory systems requires understanding this feedback cycle between the statistics of sensory signals, neural tuning properties and behavior.

The *active efficient coding* (AEC) approach offered here extends classic theories of efficient coding by a behavior component to study this feedback cycle in detail. Here we have focused on active binocular vision, where a simulated agent autonomously learns to fixate a target object with both eyes via vergence eye movements. All parts of our model self-organize in tandem to optimize overall coding efficiency. We have shown that that our model can autonomously self-calibrate and even perform accurate vergence on random dot stereograms, despite having never been exposed to such stimuli. In addition, we have reproduced various animal studies on alternate rearing conditions, which often show dramatic effects on neural representations and behavior. Our simulation results are in qualitative agreement with experimental findings, lending additional support to our model. Beyond explaining a range of experimental findings, our model also predicts systematic changes in the learned vergence behavior in response to altered regarding conditions. In addition, the model predicts that the learning of accurate vergence behavior systematically influences the neural representation and offers a novel explanation for why vertical orientations tend to be over-represented in visual cortex compared to horizontal ones, at least in primates (***De Valois et al., 1982b***) and humans (***Yacoub et al., 2008***; ***Sun et al., 2012***). These predictions should be tested in future experiments. For example, in enucleated animals, a bias in favor of vertical orientations over horizontal ones may be reduced or completely absent (***Fregnac et al., 1981***).

By freezing the neural network after the training period, we also simulated the state of the brain after the critical period. Even after fixing the optical aberrations present during training we observed a reduced vergence performance for all alternate rearing regimes. This finding is in line with a large body of evidence suggesting that optical aberrations should be corrected as early as possible to facilitate healthy development of binocular vision (e.g. ***Daw (1998***); ***Fawcett et al. (2005***), but also see ***Ding and Levi (2011***)).

While our results qualitatively match experimental findings, there are some quantitative differences. In particular, while the distribution of binocularity indices (***Wiesel and Hubel, 1963***) and disparities (***Sprague et al., 2015***) in healthy animals are relatively broad (***De Valois et al., 1982a***; ***Stevenson et al., 1992***; ***Ringach et al., 1997***), we find more narrow ones in our model. These differences are likely due to a number of simplifications present in our model. In the brain, inputs from both eyes into primary visual cortex are organized into ocular dominance bands such that individual cortical neurons may receive input which is already biased towards one or the other eye (***Le Vay et al., 1980***; ***Crowley and Katz, 2000***). In contrast, in our model all neurons receive similar amounts of input from both eyes and are therefore already predisposed for becoming binocular cells. This might explain the model’s narrower distribution of binocularity indices. Regarding the distribution of preferred disparities, animals raised under natural conditions will experience a broad range of disparities in different parts of the visual field, since objects in different locations will be at different distances. In the model, the visual input is quite impoverished, as it is dominated by a single large frontoparallel textured plane. Once this plane is accurately fixated, most parts of the visual field will appear at close to zero disparity. This may explain the narrower distribution of preferred disparities observed in the model.

Similarly, the distribution of preferred orientations in our model shows a very strong preference for horizontal and vertical, that is accentuated relative to the normal oblique effect (***Li et al., 2003***; ***De Valois et al., 1982b***). Possible reasons for this include the discrete, rectangular pixel grid with which visual inputs are sampled, the choice of our image data base (***Olmos and Kingdom, 2004a***), which contains mostly man-made structures including buildings, etc., for which it is known that they contain an abundance of horizontal and vertical edges (***Coppola et al., 1998***), and the model’s restriction to the central portion of the visual field, where the oblique effect is more pronounced (***Rothkopf et al., 2009***).

Next to addressing the above limitations, an interesting topic for future work is to use the model to study the development of amblyopia. For this, we have recently incorporated an interocular suppression mechanism, since suppression is considered a central mechanism in the development of amblyopia (***Eckmann et al., 2019***). Such models could be a useful tool for predicting the effectiveness of novel treatment methods (***Papageorgiou et al., 2019***; ***Gopal et al., 2019***).

In conclusion, we have presented a computational model that sheds new light on the central role of behavior in the development of binocular vision. The model highlights how stimulus statistics, sensory representation and behavior are all inter-dependent and influence one another and how alternate rearing conditions affect every aspect of this system. The Active Efficient Coding approach pursued here may be suitable for studying various other sensory modalities across species.

## Methods

The following paragraphs will describe the different stages of the model, the experimental setup, and the analysis. The implementation is publicly available at https://github.com/Klimmasch/AEC/.

### Image processing

We use OpenEyeSim (***Priamikov and Triesch, 2014***; ***Priamikov et al., 2016***) to render the left and right eye image. It comprises a detailed biomechanical model of the human oculomotor system and simulates a 3-dimensional environment. A rectangular plane is moved in front of the learning agent (perpendicular to the gaze direction). On it we apply greyscale textures from the McGill Calibrated Color Image Database (***Olmos and Kingdom, 2004b***) to simulate objects at different depths.

The two monocular images rendered by OpenEyeSim cover a horizontal field of view of 50° and have 320 px × 240 px (focal length *F* = 257.34 px). We use Matlab to extract single patches in different resolutions and combine corresponding patches from the left and right image. These binocular patches will be jointly encoded by the sparse coder. The *coarse scale* corresponds to 128 px× 128 px in the original image (corresponds to 26.6° × 26.6°) and is down-sampled by a factor of 4 to 32 px × 32 px. The *fine scale* image corresponds to 40 px × 40 px (8.3° × 8.3°) and is not down-sampled. From coarse and fine scale we extract 8px × 8px patches with a stride of 4 px and combine corresponding left and right patches to 16 px × 8px binocular patches (see Fig. 1). One patch in the coarse scale covers a visual angle of 6.6° and in the fine scale one patch covers 1.6°. In total, we generate 81 fine scale and 49 coarse scale patches that are subsequently normalized to have zero mean and unit norm.

### Sparse coding

The patches from coarse and fine scale are used in the sparse coding step to construct a neural representation of the visual input and to generate a reward signal that indicates the efficiency of this encoding. Each scale *S* ∈ {*c, f* } comprises a dictionary of binocular basis functions (BF) *ϕ*_*S,i*_ ∈ ℬ_*S*_ with |ℬ_*S*_| = 400. Each patch *p*_*S,j*_ 477 is reconstructed by a sparse linear combination of 10 BF:

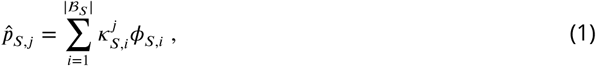

where the vector of activations 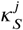 is allowed to have only 10 non-zero entries. The 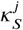 are chosen by *matching pursuit* (***Mallat and Zhang, 1993***). This greedy algorithm selects the 10 BF from the respective dictionary that yield the best approximation 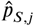 of a patch.

The reconstruction error *E*_*S*_ is calculated as the sum over all squared differences between all patches and their approximations, normalized by the total energy in the input patches:

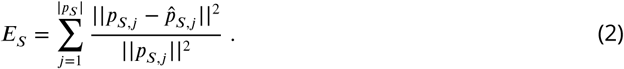

The total reconstruction error *E* = *E*_*c*_ + *E*_*f*_ is used as the negative reward (see following section) while the errors for each scale are used to update the BF (***Olshausen et al., 1996***).

The state representation is given by a feature vector, where every entry describes the mean squared activation of one BF over the whole input image:

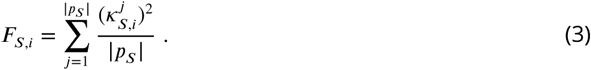

Taken together, this feature vector *F* has 2 |ℬ_*S*_| entries for both scales combined.

### Generation of motor commands

The angular position of the eyes are controlled by two extra-ocular eye muscles responsible for rotations around the vertical axis. This *medial* and *lateral rectus* are simulated utilizing an elaborate muscle model (***Umberger et al., 2003***) inside OpenEyeSim (***Priamikov and Triesch, 2014***; ***Priamikov et al., 2016***). Since we are interested in vergence movements only, we assume symmetrical eye movements so that the activities of the two muscles are mirrored for both eyes.

To generate those activations (between [0, 1] in arbitrary units) we use reinforcement learning (***Sutton and Barto, 1998***). Specifically, the model employs the CACLA+VAR algorithm from ***Van Hasselt and Wiering (2007***) that generates outputs in continuous action space. In short, it uses an actor-critic architecture (***Grondman et al., 2012***), where the actor and critic use neural networks as function approximators. These neural networks receive the state vector *S*_*t*_ that is the concatenation of the BF activations from both scales (see previous section) and the current muscle innervations. The entries in *S*_*t*_ are scaled by Welford’s algorithm (***Welford, 1962***) to have zero mean and a fixed standard deviation (0.02 in our simulations).

The critic is a one-layer network that aims to learn the value of a state. From the state vector it approximates the discounted sum of all future rewards

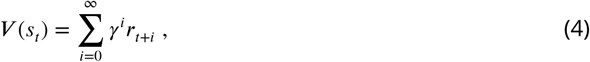

where *r*_*t*_ represents the reward achieved at time *t* and *γ* is the discount factor. To update this value network, we calculate the *Temporal Difference Error* (***Tesauro, 1995***; ***Sutton and Barto, 1998***) as *δ*_*t*_ = *r*_*t*_ + *γV*_*t*_(*S*_*t*+1_)− *V*_*t*_(*S*_*t*_). The parameters of the critic, *θ* _*V*_, are updated by

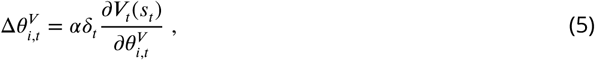

where *α* represents the learning rate for updating the critic.

The actor is a two layer artificial neural network with 50 hidden units (with tanh activation functions) and a two-dimensional output that depicts changes in muscle innervation for the two relevant eye muscles (lateral and medial rectus). The generated motor outputs are random in the beginning and the network is updated whenever the given reward was higher than estimated by the critic:

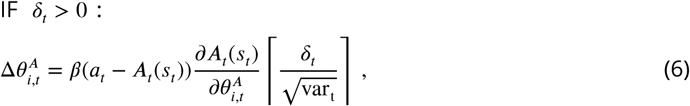

where *β* is the actor’s learning rate, *A*_*t*_(*S*_*t*_) is the action selected by the actor at time *t*, and *a*_*t*_ = *A*_*t*_ (*S*_*t*_)+ 𝒩(0, *σ*^2^) is the action that is actually executed. Adding Gaussian noise to the actor’s output to discover more favourable actions is called *Gaussian exploration*. The last term scales the update depending on how much better the action was than expected with respect to its standard deviation.

### Simulation of alternate rearing conditions

The deprivation of oriented edges is simulated by convolving the input images with elongated Gaussian kernels defined by:

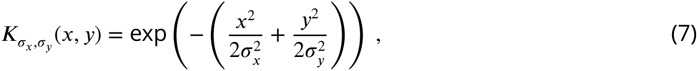

where *σ*_*x*/*y*_ represent the standard deviation in the horizontal/vertical direction.

Kernels with a large *σ*_*x*_ (*σ*_*y*_) will blur out vertical (horizontal) edges. Specifically, to simulate the deprivation of horizontal orientations, *σ*_*x*_ is set to 33 px (to cover one patch in the coarse scale completely) and *σ*_*y*_ to a small value of 0.1px. The numbers are reversed for the deprivation of vertical orientations. In the case of orthogonal rearing, the left eye receives an image deprived of horizontal orientations while the right eye receives one without vertical orientations. To make up for the small standard deviation of 0.1 in the direction that should not be impaired, the images in the *normal* case are convolved with a Gaussian kernel with *σ*_*x*_ = *σ*_*y*_ = 0.1px.

To simulate monocular deprivation (MD) we set *σ*_*x*_ = *σ*_*y*_ = 240 px for the right input image only. The small patches that we extract from this strongly blurred image contain almost no high spatial frequencies.

A strabismus is artificially induced by rotating the right eye ball inwards as it is commonly done in biological experiments by fixating a prism in front of the eye or by cutting the lateral rectus muscle. In our Open-Eye-Simulator, however, we can rotate the eye by a specific angle. One input patch in the coarse scale covers 6.6°. When we set the strabismic angle to 3° there is still an overlap in the input images that will be reflected in the neural code. In contrast, when the strabismic angle is set to 10°, the input patches become completely uncorrelated. Examples of the changes done to the input images are displayed in Fig. 2.

### Analysis of receptive fields

To determine the orientations of the basis functions (BFs) we use MATLAB’s implementation of the *trust region reflective algorithm* for non-linear curve fitting (***Coleman and Li (1996***)) to fit them to two-dimensional Gabor functions as defined by:

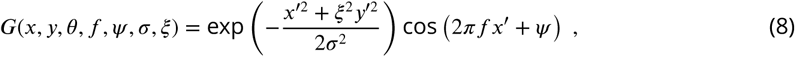

with *x*′ = *x* cos (*θ*) + *y* sin (*θ*) and *y*′ = −*x* sin (*θ*) + *y* cos (*θ*).

Here, *f* denotes the frequency, *Ψ* the phase offset, *σ* the standard deviation of the Gaussian envelope, *ξ* the spatial aspect ratio and *θ* the orientation, where *θ* =0 deg corresponds to a vertically oriented Gabor function. We initialize the parameters randomly 150 times and fit the function either to the left or right BFs (or to both, see below). To evaluate the quality of the fits, we record the difference between the actual BFs and the Gabor fit. More specifically, the *residual* is defined as the sum of the squared differences in single pixel values between BFs and the fit. To compare the fits across the different experimental conditions, we only took those fits where this residual was less than or equal to 0.2. This accounts for more than 96% of all BFs in the healthy case.

Another interpretation for these fits is a stimulus that maximally activates the particular neuron. To investigate the binocularity of such a cell, we compare their monocular response to the left and right Gabor fit. The eye with the greater response is the dominant eye for this neuron. Similar as in ***Hubel and Wiesel (1962***) we show the best stimulus (here the Gabor fit) to the dominant eye and the same stimulus to the non-dominant eye and record the responses *L* and *R*. We then compare these by

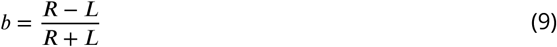

to get a binocularity index between -1 (monocular left) and +1 (monocular right), where 0 means perfectly binocular.

When fitting this function to binocular BFs, we assume that all parameters are equal for the left and right monocular sub-region of the BFs except for the phase offset *Ψ*, that can be different for left and right eye. Following the assumption that the disparity tuning in a binocular cell is encoded by means of this phase shift, we can calculate the preferred disparity *d* of a neuron by:

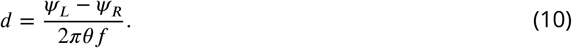

The maximally detectable disparity is given by the RF size, that is, the visual angle one binocular patch covers. BFs with a disparity preference bigger than that are excluded from the analysis.

### Laplacian disparity distribution

The probability density distribution of a Laplacian distributed random variable *X* is defined as

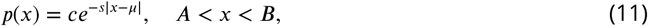

To simulate the disparity distribution we set *μ* to the angle that is desired to fixate an object at a certain distance *d*_*o*_

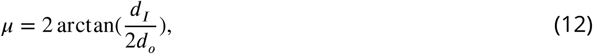

where *d*_*I*_ = 56 cm is the interpupillary distance. The data shown in Fig. 8B are generated from a model with only the fine scale, for simplicity.

## Acknowledgments

This work was supported by the German Federal Ministry of Education and Research under Grants 01GQ1414 and 01EW1603A (within the frame of ERA-NET NEURON), the European Union’s Horizon 2020 Grant 713010, the Hong Kong Research Grants Council under Grant 16244416, and the Quandt Foundation.

## Appendix 1

**Appendix 1 Figure 1.**
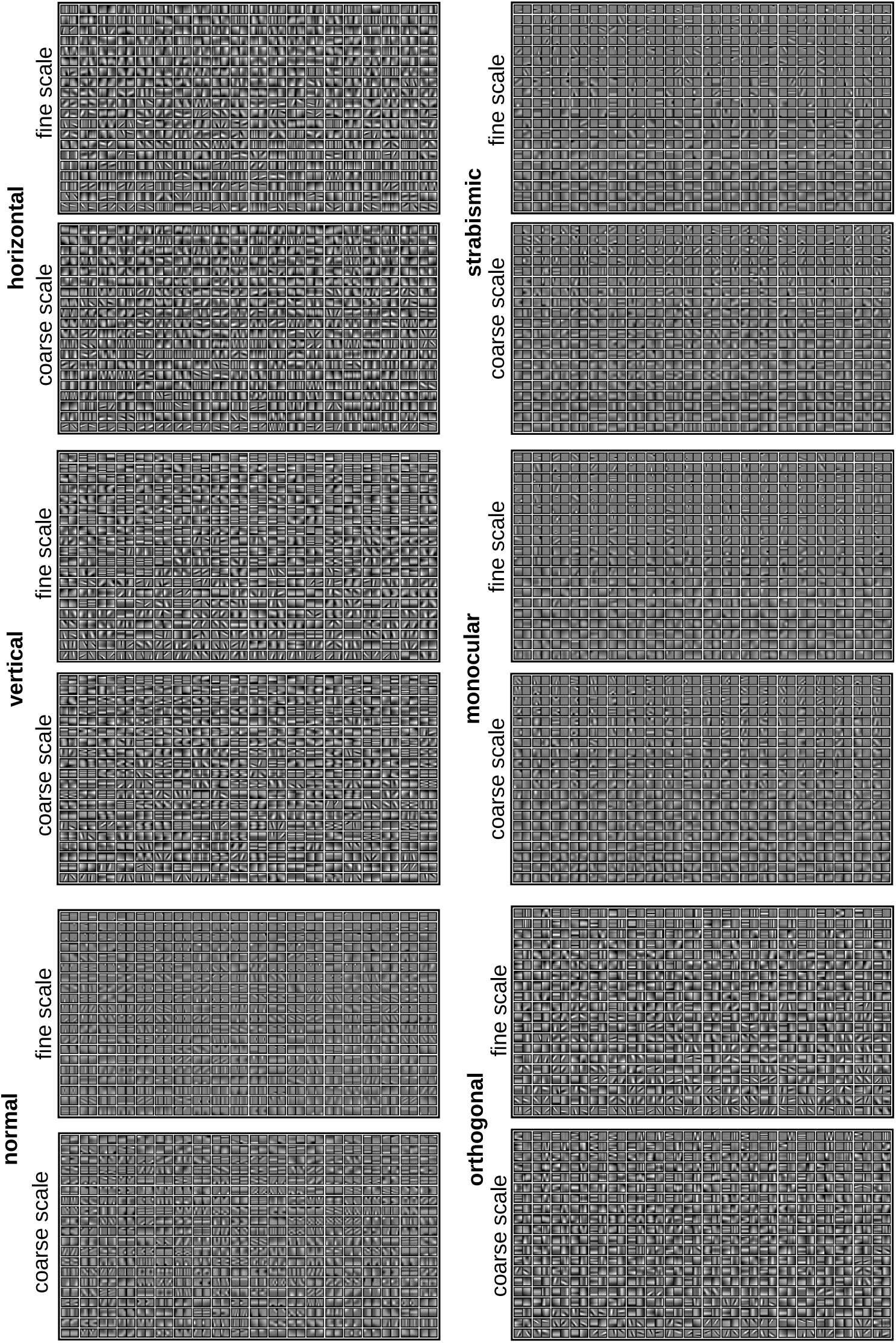
Complete set of all BFs that are learned during training for all different rearing conditions.

## Appendix 2

**Appendix 2 Figure 1.**
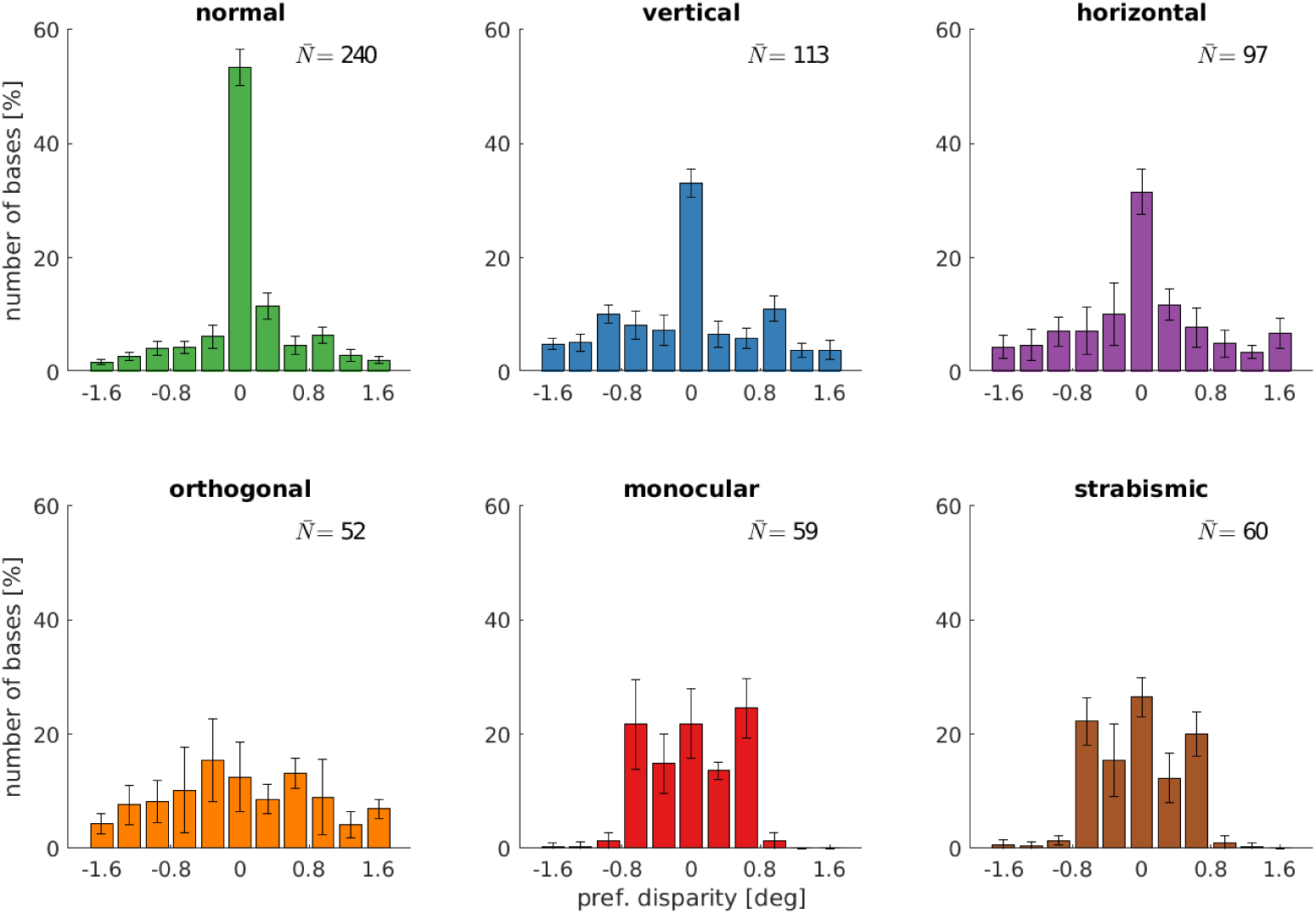
Disparity tuning of the fine scale of models that were trained under different rearing conditions.

## Appendix 3

**Appendix 3 Figure 1.**
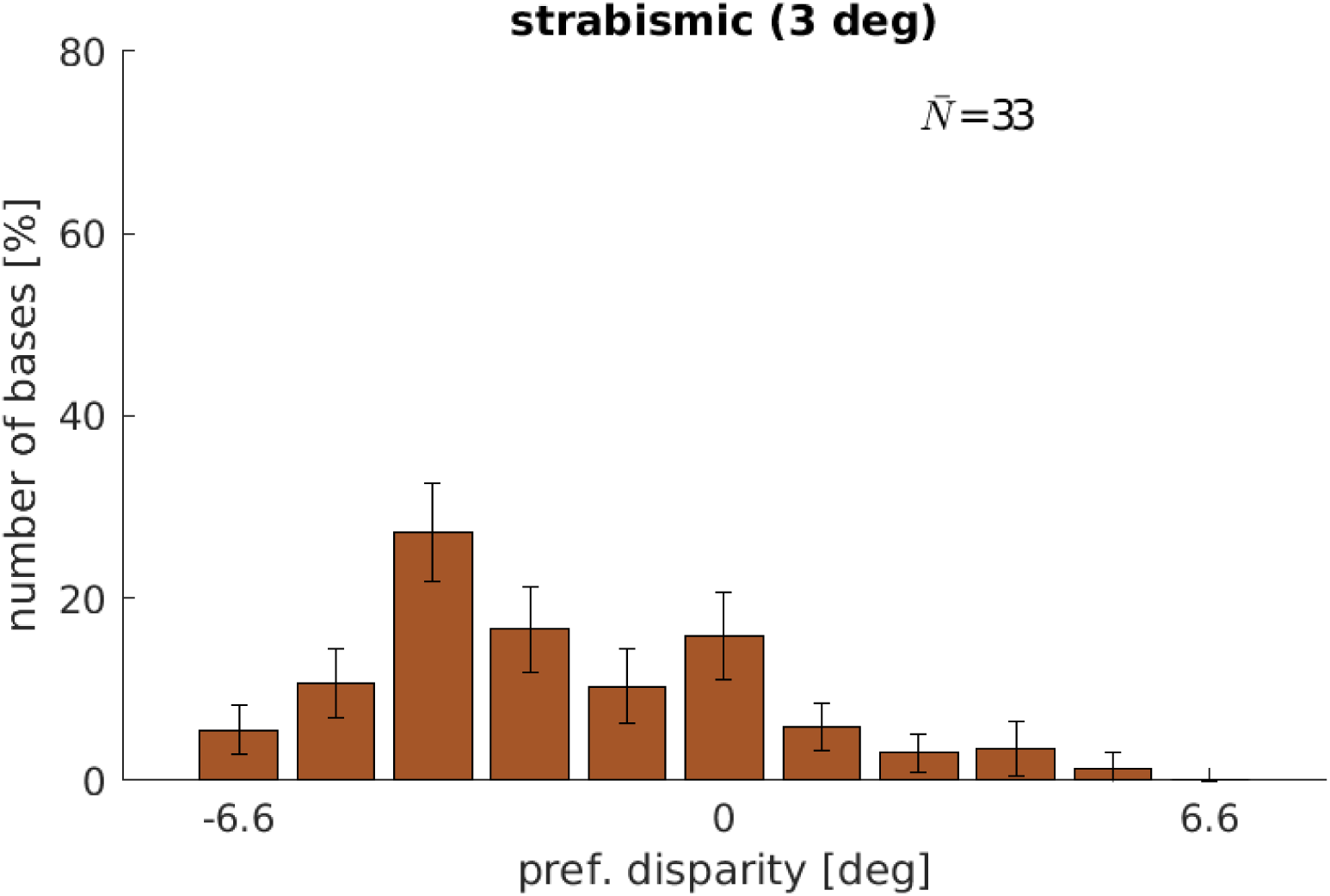
Disparity tuning of a model that was trained with a constant strabismic angle of 3 deg. Note the marked similarity to ***Shlaer (1971***), Fig. 2. Depicted are the results from the coarse scale only.

